# Dynamic cortical and tractography atlases of proactive and reactive alpha and high-gamma activities

**DOI:** 10.1101/2022.07.16.500323

**Authors:** Hiroya Ono, Masaki Sonoda, Kazuki Sakakura, Yu Kitazawa, Takumi Mitsuhashi, Ethan Firestone, Aimee F. Luat, Neena I. Marupudi, Sandeep Sood, Eishi Asano

## Abstract

Alpha waves - posterior-dominant rhythms at 8-12 Hz reactive to eye opening and closure - are among the most fundamental EEG findings in clinical practice and research since Hans Berger first documented them in the early 20th century. Yet, the exact network dynamics of alpha waves in regard to eye movements remains unknown. High-gamma activity at 70-110 Hz is also reactive to eye movements and a summary measure of local cortical activation supporting sensorimotor or cognitive function. We aimed to build the first-ever brain atlases directly visualizing the network dynamics of eye movement-related alpha and high-gamma modulations, at cortical and white matter levels. We studied 28 patients (age: 5-20 years) who underwent intracranial EEG and electrooculography recordings. We measured alpha and high-gamma modulations at 2,170 electrode sites outside the seizure onset zone, interictal spike-generating areas, and MRI-visible structural lesions. Dynamic tractography animated white matter streamlines modulated significantly and simultaneously beyond chance, on a millisecond scale. *<u>Before eye closure onset</u>*, significant alpha augmentation occurred at the occipital and frontal cortices. *<u>After eye closure onset</u>*, alpha-based functional connectivity was strengthened, while high gamma-based connectivity was weakened extensively in both intrahemispheric and interhemispheric pathways involving the central visual areas. The inferior fronto-occipital fasciculus supported the strengthened alpha coaugmentation-based functional connectivity between occipital and frontal lobe regions, whereas the posterior corpus callosum supported the interhemispheric functional connectivity between the occipital lobes. *<u>After eye opening offset</u>*, significant high gamma augmentation and alpha attenuation occurred at occipital, fusiform, and inferior parietal cortices. High gamma coaugmentation-based functional connectivity was strengthened, whereas alpha-based connectivity was weakened in the posterior interhemispheric and intrahemispheric white matter pathways involving central and peripheral visual areas. *<u>Proactive</u>* and *<u>reactive</u>* alpha waves involve extensive, distinct white matter networks that include the frontal lobe cortices, along with low- and high-order visual areas. High-gamma co-attenuation coupled to alpha co-augmentation in shared brain circuitry after eye closure supports the notion of an idling role for alpha waves during eye closure. These dynamic tractography atlases may improve understanding of the significance of EEG alpha waves in assessing the functional integrity of brain networks in clinical practice; they also may help elucidate the effects of eye movements on task-related brain network measures observed in cognitive neuroscience research.

## Introduction

Our primary goal for this study was to build a large-scale human brain atlas animating the rapid cortical dynamics and white matter functional connectivity of alpha waves; these brain oscillations are posterior-dominant rhythms at 8-12 Hz modulated by eye closure and opening. For the first time in 1929, Hans Berger used scalp electroencephalography (EEG) to document alpha waves, which were distributed mainly in the bilateral posterior head regions, augmented with eye closure, and attenuated with eye opening.^1^ Since their discovery, eye-movement reactive alpha waves have been among the most fundamental EEG findings in clinical practice and research, and the absence of reactive alpha waves suggests functional impairment of a given hemisphere.^2-9^ However, alteration of reactive alpha waves on scalp EEG may not effectively localize the pathological brain networks, perhaps because the sources and circuitry are poorly understood.^10^ Existing evidence from scalp EEG and magnetoencephalography (MEG) studies suggest that reactive alpha waves primarily originate from the lower-order visual cortex because their amplitude is highest in occipital areas and modulated by visual stimuli.^2,11-13^ Yet, studies using noninvasive recordings may not have sufficient signal fidelity^14^ to clarify the spatiotemporal dynamics of eye-movement-related alpha waves, as they propagate through the inferior or medial surfaces of the cerebral cortex.

Power,^15-17^ coherence,^18^ and traveling waves^19,20^ on invasive recordings are valuable measures to evaluate the network dynamics of reactive alpha waves. Coherence and traveling waves quantify connectivity strength based on oscillatory phase relationships between electrode sites, whereas power denotes the intensity of EEG signals. A study in dogs demonstrated that alpha wave power was increased in the lower-order visual cortex following eye closure, and a phase reversal of such reactive alpha waves was noted at 1.1 mm below the pia mater within the gray matter.^21^ Another study in dogs demonstrated that coherence-based connectivity of reactive alpha waves was more robust between distant visual cortices than between the visual cortex and the thalamic lateral geniculate nucleus (LGN).^18^ Granted, other investigations have demonstrated that reactive alpha waves co-occurred in the visual cortex and thalamus.^10,22^ These findings suggest that the origins of reactive alpha waves include but are not necessarily confined to the lower-order visual cortex.

Presurgical evaluation of patients with drug-resistant focal epilepsy has provided a unique opportunity to record human intracranial EEG (iEEG) signals directly from the occipital areas during wakefulness. A previous iEEG study of seven patients with focal epilepsy identified alpha waves during eye closure traveling from the temporal-parietal areas toward the lateral occipital and anterior temporal regions.^23^ Another iEEG study of five patients reported that alpha waves during eye closure traveled from higher- to lower-order visual areas and that alpha waves in the cortex preceded those in the thalamic pulvinar nucleus; the investigators inferred that observed alpha waves might reflect feedback rhythms propagating from higher to lower-order visual cortex.^24^ Traveling wave analysis is designed to only determine the direction of *<u>local</u>* neural propagations based on the phase of sustained oscillations; thus, the traveling wave-based studies mentioned above^23,24^ could only characterize alpha wave propagations within the several centimeters of each grid or strip electrode array. Therefore, the degree of alpha-based connectivity outside the visual areas remains poorly understood.

The spatial extent of signal sampling is inevitably limited in any iEEG study of the human brain because electrode coverage is strictly decided by clinical necessity. To overcome this issue, one needs to analyze iEEG signals sampled from multiple patients using a method agnostic to the oscillatory phase information. Thus, the present study quantified iEEG alpha amplitude (i.e., square root of power) time-locked to eye movements at 2,170 non-epileptic electrode sites sampled from 28 patients. We then animated the spatio-temporal dynamics of eye movement-related alpha amplitude augmentation and attenuation on a spatially normalized brain template.^25-27^ **[Aim 1]** We determined whether alpha amplitude augmentation following eye closure would take place initially at lower-order and subsequently at higher-order visual areas (i.e., in a feedforward direction) or in the opposite (feedback) direction. We likewise characterized the spatio-temporal dynamics of alpha amplitude attenuation following eye opening.

The present study aimed to signify the functional roles of eye movement-related alpha modulations by correlating them with high-gamma activity at 70-110 Hz at a given space and time. Extensive evidence suggests that high-gamma amplitude augmentation reflects local cortical activation. For example, it is associated with increased neural firing,^28-30^ glucose metabolism on positron emission tomography,^31^ and hemodynamic activation on functional magnetic resonance imaging (fMRI).^16,28,32,33^ In contrast, high-gamma attenuation was associated with reduced neural firing and hemodynamic deactivation.^34^ A meta-analysis of 15 iEEG studies suggests that task-related high-gamma augmentation can accurately localize the eloquent cortices defined by electrical stimulation mapping.^35^ Our previous iEEG study of 65 patients with drug-resistant focal epilepsy demonstrated that language task-related high-gamma activity predicted postoperative neuropsychological impairment with an accuracy of 0.80.^36^ Investigators have proposed that alpha waves augmented after eye closure, at least in part, exert functional idling or inhibition on cortical areas.^2,3,37^ We expected that our intracranial EEG study would provide evidence supporting such a role. **[Aim 2]** We tested the hypothesis that cortical networks showing alpha augmentation after eye closure would simultaneously exhibit high-gamma attenuation. We likewise tested the hypothesis that those showing alpha attenuation after eye opening would simultaneously exhibit high-gamma augmentation.

The present study aimed to animate the dynamics of intra- and inter-hemispheric functional connectivity pathways by combining the timing of eye movement-related iEEG amplitude modulations at various sites with diffusion-weighted imaging (DWI) white matter tractography. A previous study in dogs demonstrated that inter-hemispheric coherence in reactive alpha waves was strong between homotopic visual cortices.^18^ Scalp EEG studies of patients with drug-resistant epilepsy reported that corpus callosotomy reduced inter-hemispheric alpha-band coherence between the occipital lobes immediately after disconnection.^38,39^ These studies provide causal evidence that homotopic visual cortices showing reactive alpha waves have functional connectivity via *<u>direct</u>* inter-hemispheric cortico-cortical white matter pathways. Other investigators have provided evidence that neuronal circuits simultaneously engaging in high-frequency activities (e.g., high-gamma co-augmentation) are susceptible to developing use-dependent, direct functional connectivity.^40-42^ Many fMRI, MEG, scalp EEG, and iEEG studies have inferred that cortices with neural or hemodynamic responses generated simultaneously and consistently across trials beyond chance are functionally connected.^4,43-52^ The novelty of our dynamic connectivity atlas in this study is visualizing direct white matter pathways connecting two remote cortical regions and assessing iEEG-based functional connectivity with a temporal resolution in the millisecond range. **[Aim 3]** We determined whether white matter functional connectivity between cortical areas simultaneously showing eye movement-related alpha modulations would initially involve the lower-order and followed by the higher-order visual networks. We likewise determined the spatio-temporal dynamics of white matter functional connectivity alteration between cortical areas showing high-gamma co-modulations.

## Materials and Methods

### Patients

We studied 28 patients with drug-resistant focal epilepsy (age range: 5-20 years; 15 females; Table 1) who satisfied the following eligibility criteria. The inclusion criteria included [a] simultaneous video-iEEG and electrooculography (EOG) recordings from December 2008 to July 2018, as part of our presurgical assessment at Children’s Hospital of Michigan, Detroit, MI,^53^ [b] iEEG sampling from the occipital lobe, and [c] at least 16 events of spontaneous eye closure and eye opening detected by both video and EOG during wakefulness (see the Methods section below). The exclusion criteria included [a] age of less than five years (because the posterior dominant rhythm may range below 8 Hz in healthy children,^54^ [b] presence of seizure onset zone (SOZ),^55^ interictal spike discharges,^56^ or MRI lesions affecting the occipital lobe, [c] visual field deficits by confrontation, [d] history of previous epilepsy surgery, and [e] posterior dominant rhythm slower than 8 Hz on a preoperative scalp EEG recording. The study was approved by the Wayne State University Institutional Review Board. Written informed consent was obtained from the patients’ legal guardians, and written assent was also collected from the patients, when possible.

**Table 1.** Patient profile. SD: standard deviation.

|  |  |
| --- | --- |
| Patient profile. |  |
| Number of patients | 28 |
| Mean age in years (range) | 12.5 (5 - 20) |
| Proportion of female (%) | 53.6 |
| Sampled hemisphere (%) |  |
| Left | 46.4 |
| Right | 53.6 |
| Seizure onset zone (%) |  |
| involving the frontal lobe | 28.6 |
| involving the temporal lobe | 39.3 |
| involving the parietal lobe | 25.0 |
| Proportion of patients with an MRI lesion (%) | 67.9 |
| Mean number of antiepileptic drugs (SD) | 1.9 (0.65) |

### iEEG and three-dimensional (3D) MRI surface imaging data acquisition

The iEEG and MRI data acquisition protocols were identical to those reported in our previous studies.^27,42^ We implanted platinum subdural electrodes (3 mm diameter and 10 mm center-to-center distance) on the affected hemisphere. The spatial extent of iEEG sampling was determined by clinical needs, and the goal of iEEG recording was to determine the boundaries between the presumed epileptogenic zone and eloquent areas. We continuously acquired video-iEEG data with a sampling rate of 1,000 Hz at the bedside, for 2-5 days following intracranial electrode placement. We did not expand the extent nor duration of iEEG recording for research purposes. We only included artifact-free, non-epileptic electrode sites: defined as those unaffected by the SOZ, interictal spikes, or MRI lesions.^57,58^ Thus, 2,170 electrode sites (mean: 77.5 per patient; range: 36 to 108) were available for the following iEEG analysis (Fig. 1A). We took this analytic approach to minimize the unwanted effects of pathological high-frequency oscillations^59,60^ on the measurement of eye movement-related iEEG responses.

**Figure 1.**
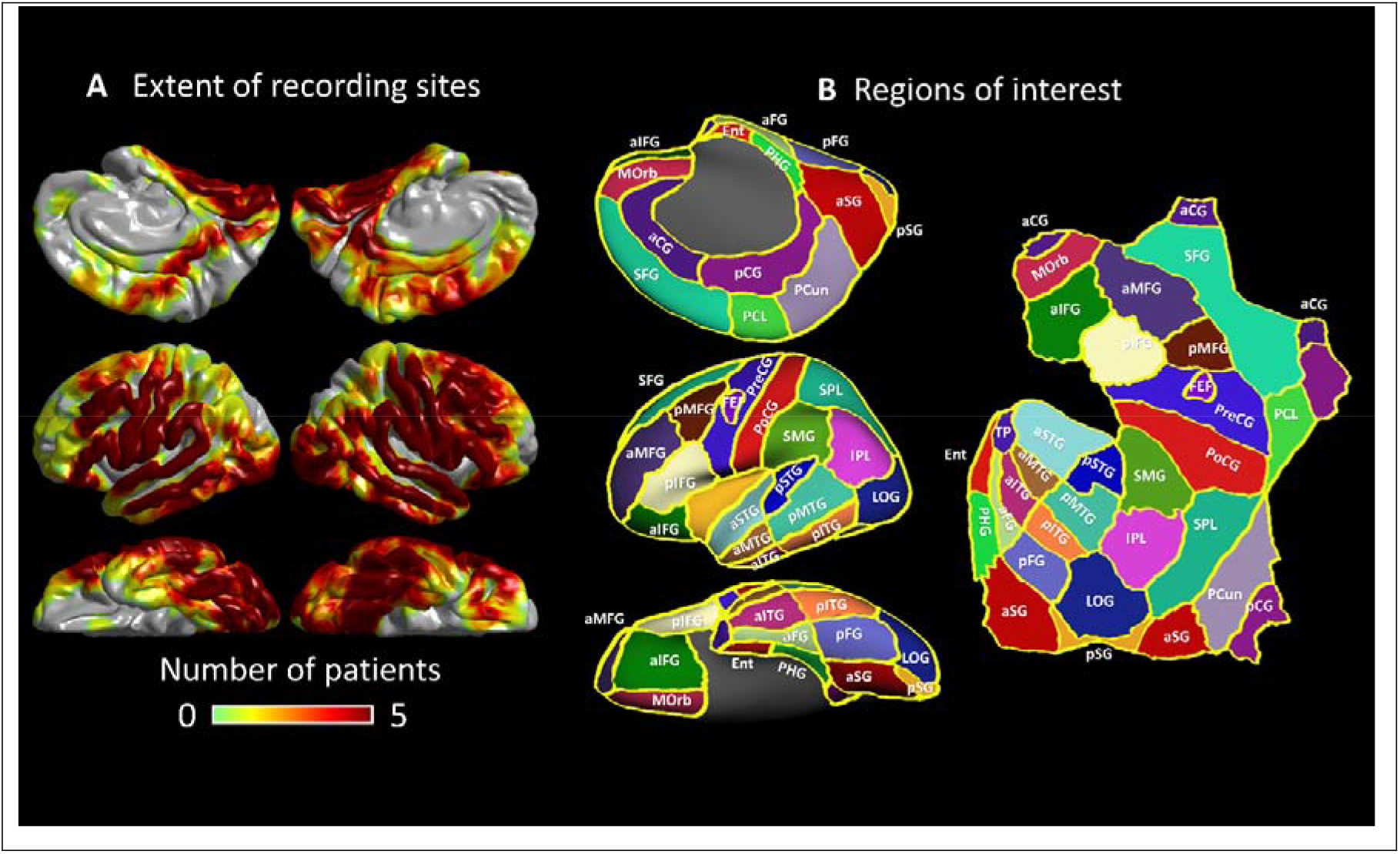
The spatial extent of non-epileptic electrode sites on a standard brain surface image. (A) Color indicates the number of patients available at a given analysis mesh. The number of non-epileptic electrode sites in each region of interest (ROI) is provided in Supplementary Table 1. All 28 patients had at least one subdural electrode site in each of the four lobes. (B) The locations of ROIs on the left hemisphere are presented. The striatal cortex consisted of the summation of lingual and cuneus gyri; thereby, the anatomical boundary between the anterior and posterior striatal cortex was defined as the line connecting ‘the boundary with the posterior fusiform and lateral occipital regions’ and ‘that with the precuneus and superior parietal lobule’. Frontal eye fields (FEFs) were defined based on the results of electrical stimulation mapping as reported in our previous studies.^7,123^ Refer to Supplementary Table 1 for the meaning of each abbreviation.

We built a 3D surface image with subdural electrodes displayed on the pial surface using a preoperative T1-weighted spoiled gradient-recalled echo sequence MRI and a CT image taken immediately after the placement of subdural electrode arrays.^27,61^ For group-level visualization and analysis of iEEG alpha and high-gamma modulations, we normalized each electrode site to the FreeSurfer standard coordinates, referred to as FSaverage (http://surfer.nmr.mgh.harvard.edu). Using the Desikan anatomical parcellation,^25,27^ we divided the cerebral cortex of each hemisphere into 30 regions of interest (ROIs) (Fig. 1B). We employed a quantitative iEEG analysis only at 52 ROIs (i.e., 26 × 2) that included at least four electrode sites on each hemisphere (see the detail in Supplementary Table 1).

### Marking of eye closure and opening events

As part of our routine presurgical evaluation, we placed EOG electrodes 2.5 cm below and 2.5 cm lateral to the left and right outer canthi and continuously monitored spontaneous eye movements during the video-iEEG recording.^62,63^ While blinded to the results of the iEEG time-frequency analysis, the first author (H.O.) visually marked the onset and offset of at least 16 events of spontaneous eye-closure and eye-opening during a task-free, wakeful resting state, under room light, using video and EOG recordings. We did not include the events of eye closure preceded or followed by opening within two seconds nor opening preceded or followed by closure within two seconds. We also did not include eye movement events not confirmed by video. Patients were not instructed to watch or attend visual stimuli for this study. We excluded iEEG periods during cognitive tasks^64^ or within two hours following seizure events. On EOG recording with a time constant of 1.0 second, we defined the onset and offset of each eye closure (Fig. 2A) or eye opening (Fig. 2B) event as the onset and offset of signal deflection greater than 50 μ V.^65^ The mean number of identified eye closure events was 35.9 per patient (range: 18-66 per patient), whereas that of eye-opening events was 34.9 per patient (range: 16-77 per patient).

**Figure 2.**
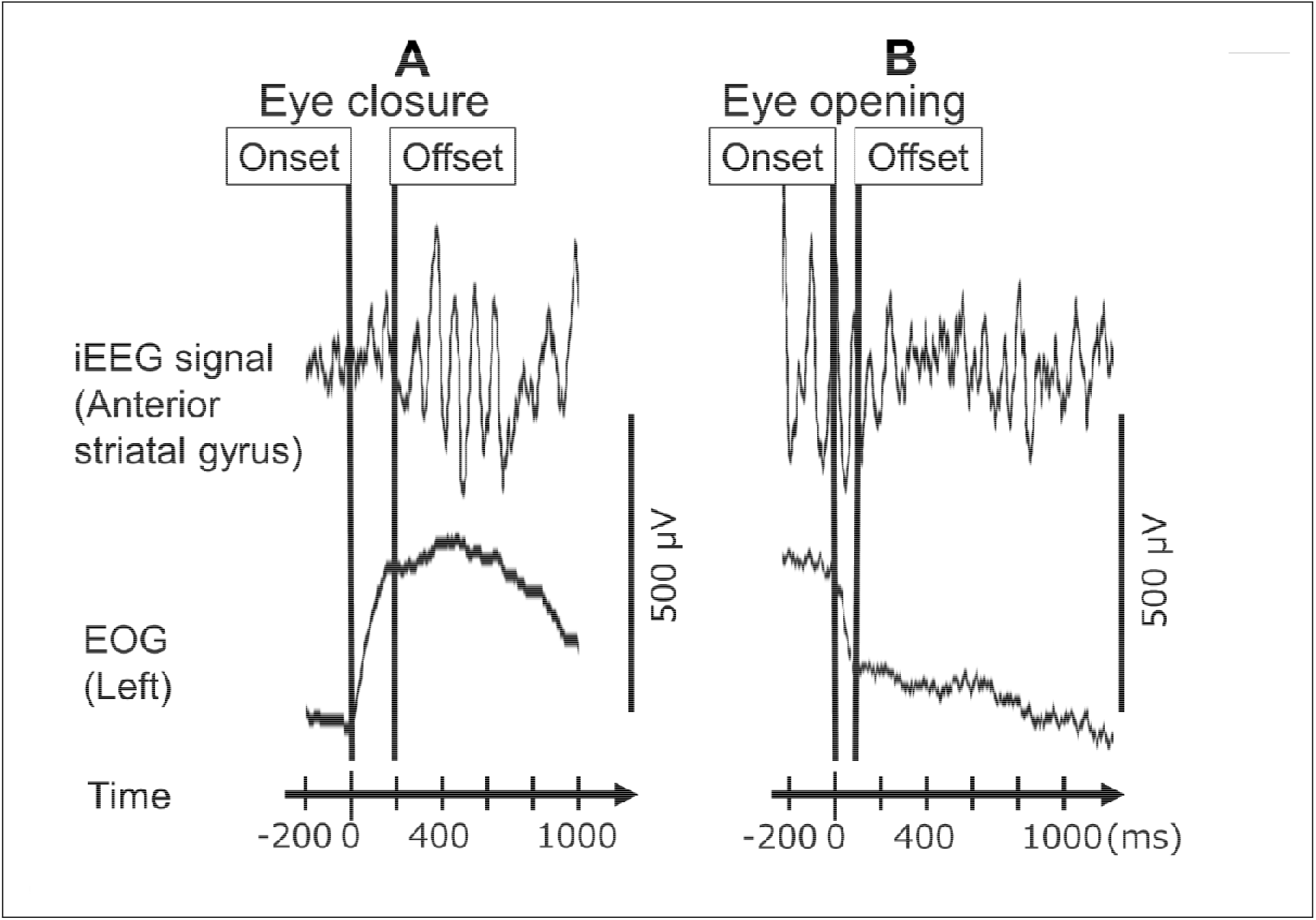
Marking of onset and offset of eye movements in a 12-year-old girl with focal epilepsy. (A) A peri-eye closure period. (B) A peri-eye opening period. Upper trace: Intracranial EEG (iEEG) signals at a non-epileptic anterior striatal gyrus (aSG) site in the occipital lobe (high-cut filter: 300 Hz. time constant: 0.1 s). Lower trace: Electrooculography (EOG) signals to determine the onset and offset of given eye movements.

### Time-frequency analysis to determine the dynamics of iEEG amplitude modulations

At each electrode site, we determined how much (% change) alpha or high-gamma amplitude (i.e., square root of power) was augmented or attenuated, at given time bins, compared to the baseline (i.e. 200 and 600 ms before the eye movement onset). To this end, we employed a complex demodulation method^66^ identical to that reported in our previous iEEG studies of sensorimotor and cognitive processing.^27,42^ This method - implemented in BESA EEG Software (BESA GmbH, Gräfelfing, Germany)^67^ - transformed time-voltage signals into time-frequency bins. The time-frequency transformation was done by multiplying the time-domain iEEG signal with a complex exponential, followed by a band-pass filter. This method was equivalent to a Gabor transformation because it employed a Gaussian-shaped low-pass finite impulse response (FIR) filter. The size of each time-frequency bin was ‘25 ms × 2 Hz’ for measuring alpha modulations (8-12 Hz); the time-frequency resolution was ±39.4 ms and ±2.8 Hz, and defined as a 50% power drop of the FIR filter. We analyzed high-gamma modulations with ‘5 ms × 10 Hz’ time-frequency bins; the time-frequency resolution was ±7.9 ms and ±14.2 Hz. We employed the time-frequency transformation for each 3,000-ms time window ranging from 1,000 ms before and 2,000 ms after each of the following behavioral marks: [i] eye-closure onset, [ii] eye-closure offset, [iii] eye-opening onset, and [iv] eye-opening offset (Fig. 2).

### Region of interest (ROI) analysis of iEEG amplitude modulations

The studentized bootstrap statistics determined when (at what time bins) and where (at what Desikan-based anatomical ROIs; Fig. 1B) eye movement-related alpha or high-gamma amplitude differed from the mean, during the baseline period between 200 and 600 ms before eye movement onset (Fig. 3 and Supplementary Video 1).^64^ We thereby considered significant events as iEEG amplitudes modulated beyond the 99.99% confidence intervals (99.99% CI) for at least eight consecutive time-bins (corresponding to the duration of 2 oscillatory cycles).^64^ Employment of a 99.99% CI is equivalent to the Bonferroni correction for 500 repeated comparisons. Such a strict threshold might increase the risk of Type II errors but reduce Type I errors, considering the total number of 25-ms and 5-ms time bins in 2,200-ms time windows (e.g., a period between 200 ms before and 2,000 ms after eye closure onset).

**Figure 3.**
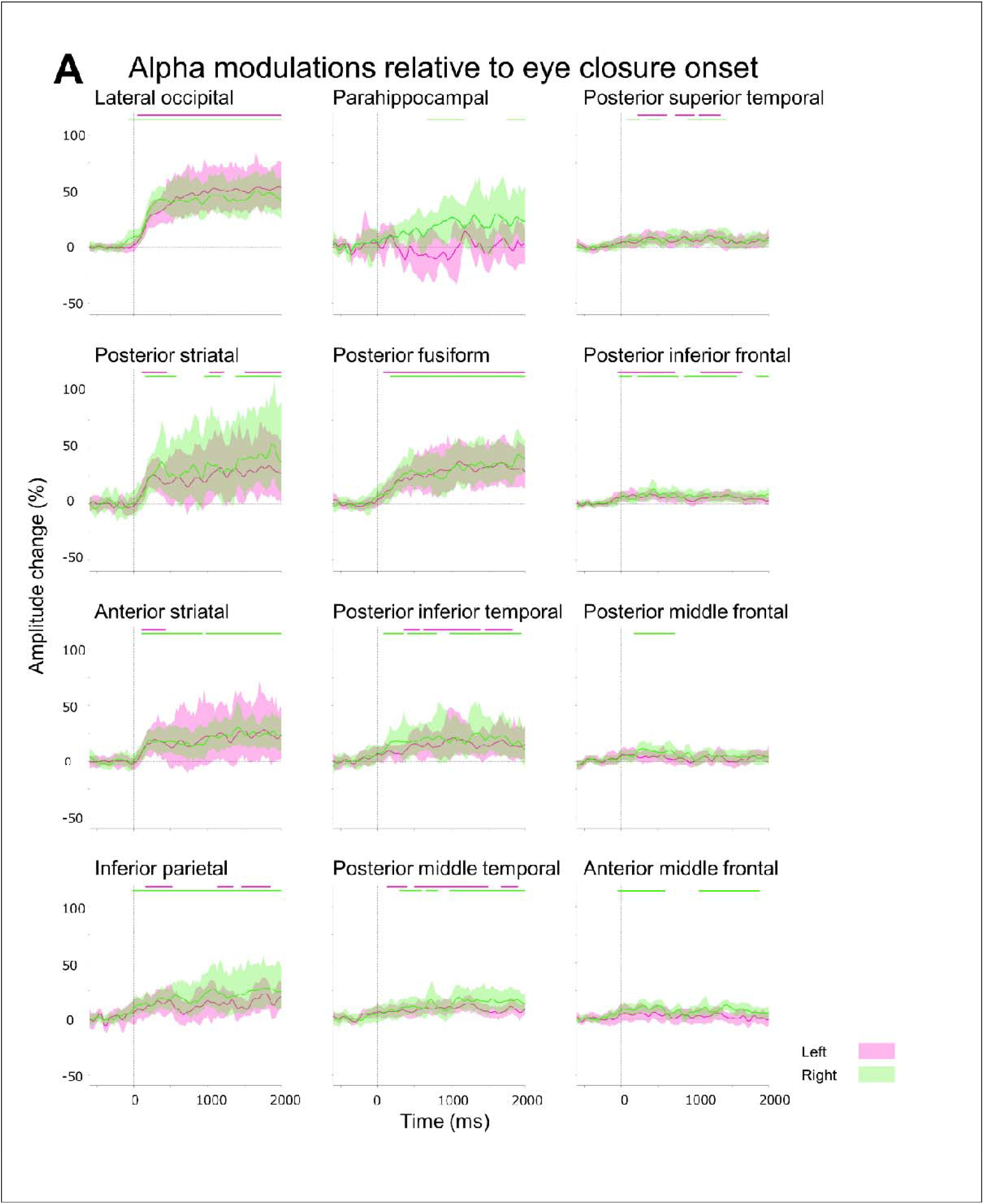
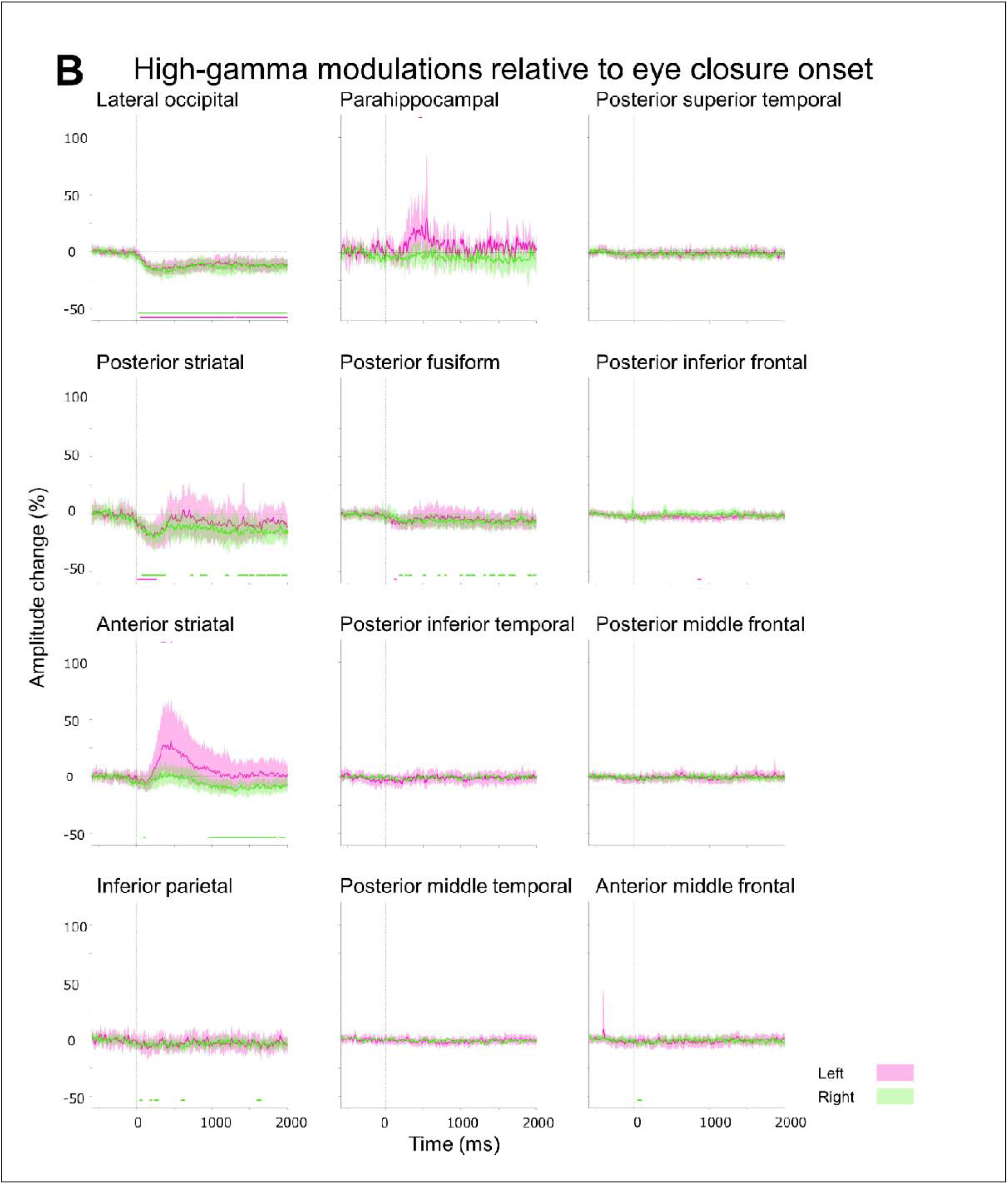
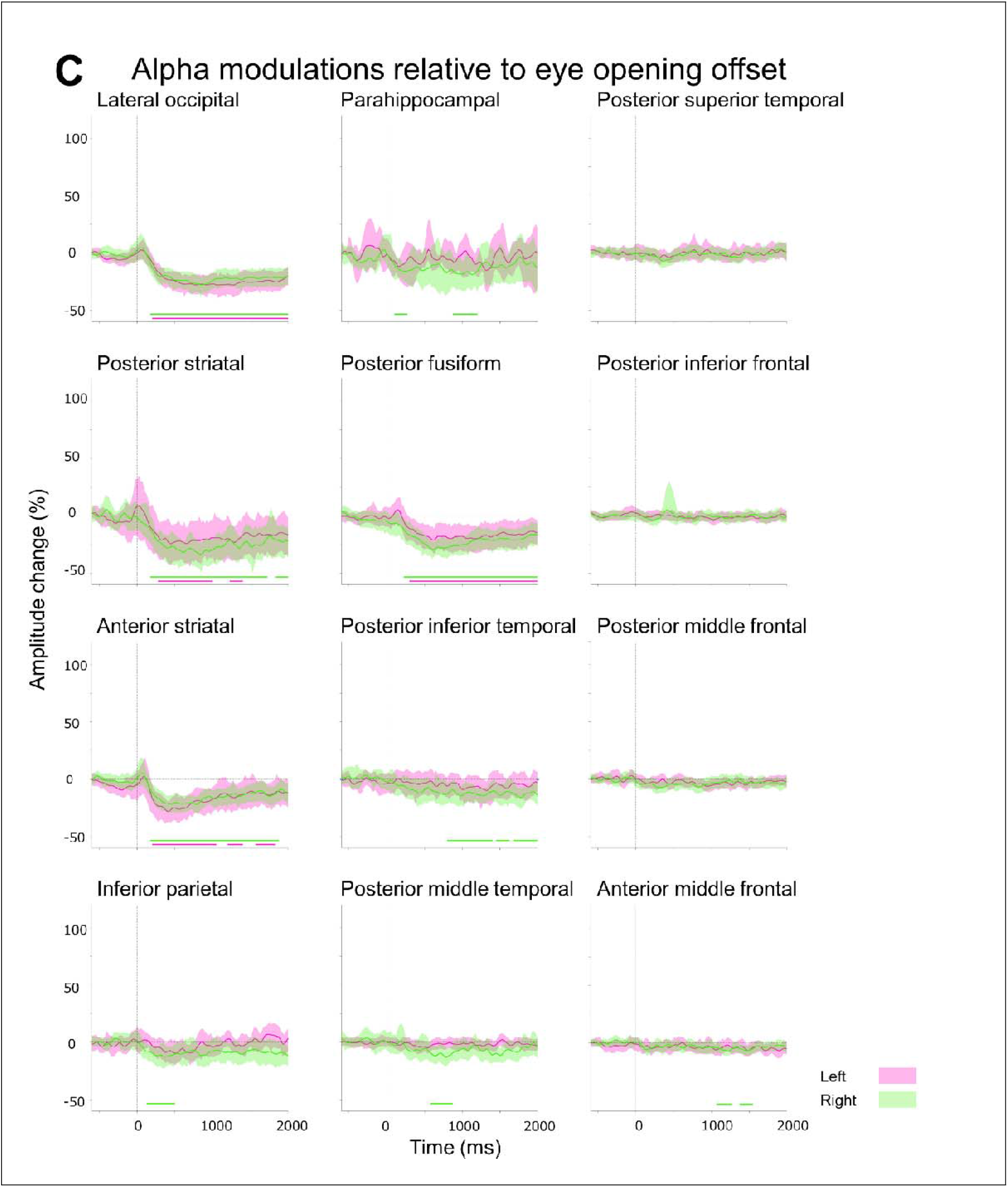
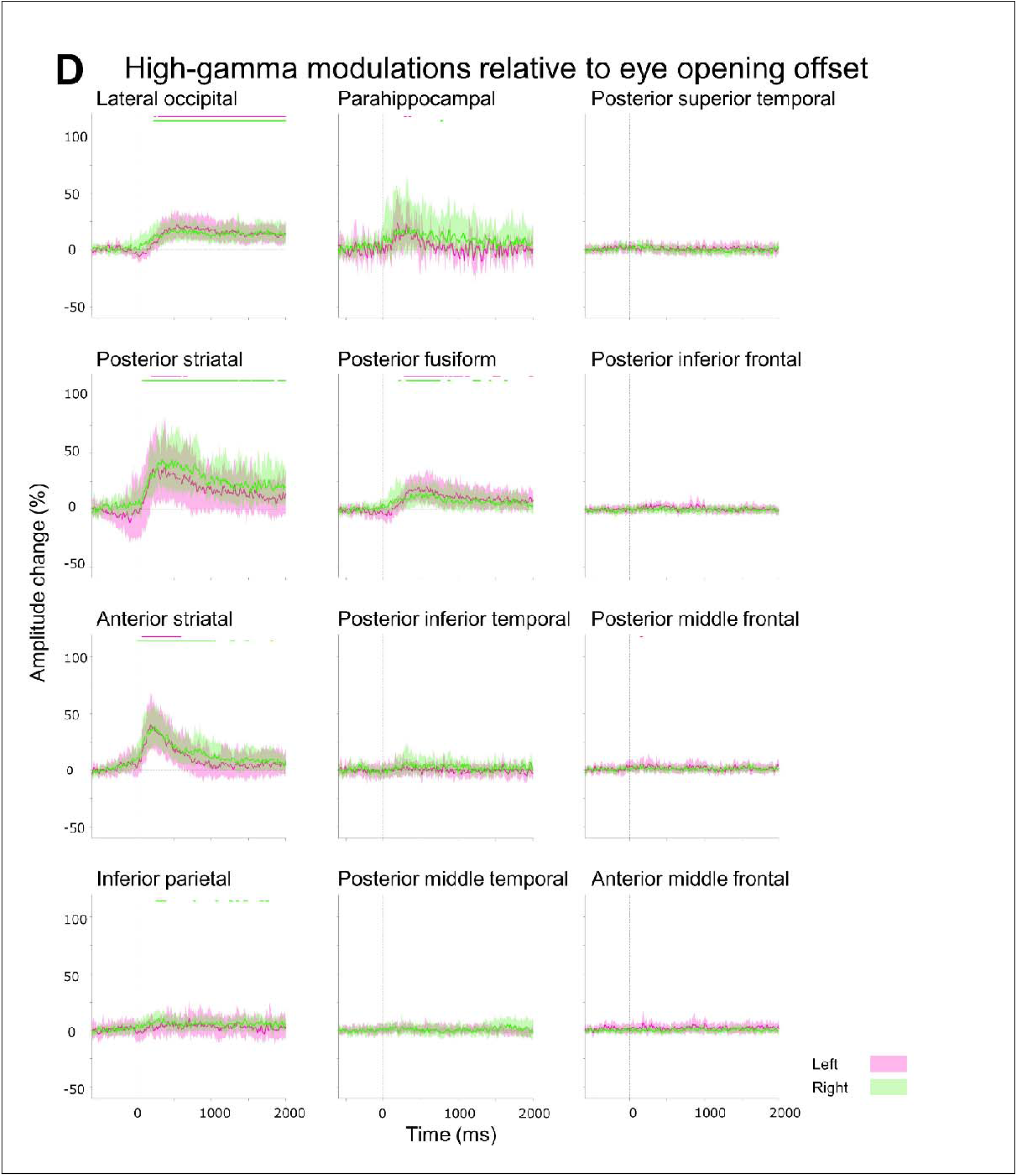
Eye movement-related alpha and high-gamma modulations at regions of interest (ROIs). A given plot presents the percent change in alpha and high-gamma amplitudes at each Desikan anatomical ROI compared to the baseline at 200 to 600-ms before eye movement onset. Solid line: mean across all available electrode sites within a given ROI. Shade: 99.99% confidence interval. Magenta: Left hemisphere. Green: Right hemisphere. (A and B) Eye closure-related alpha and high gamma modulations (0 ms: eye closure onset). (C and D) Eye opening-related alpha and high gamma modulations (0 ms: eye opening offset). Supplementary Video 1 presents eye movement-related alpha and high-gamma modulations at all analyzed ROIs.

[**Aim 1**] The aforementioned ROI analysis (Fig. 3A) effectively determined the temporal order of significant eye closure-related alpha augmentation across the lower- and higher-order visual areas, as well as non-visual areas. In the present study, the lower-order visual areas included the posterior striatal, anterior striatal, and lateral occipital regions. In contrast, the higher-order visual areas included the parahippocampal, posterior fusiform, posterior inferior temporal gyrus (pITG), and inferior parietal gyri.^68-71^

[**Aim 2**] We tested the hypothesis that eye closure-related alpha augmentation co-occurred with high-gamma attenuation, at given anatomical ROIs (Fig. 3A and B). To this end, we computed ‘alpha-by-high-gamma multiplied value’ defined as a multiplication of alpha and high-gamma amplitude (% change) at each of the eighty-nine 25-ms epochs between -200 ms before eye closure onset and +2,000 ms for each ROI. A one-sample t-test determined which ROIs had the mean alpha-by-high-gamma multiplied value differing from zero. If this value was below zero, our hypothesis would be supported. We employed a Bonferroni correction for 52 ROIs. Likewise, we tested the hypothesis that eye opening-related alpha attenuation co-occurred with high-gamma augmentation at given ROIs (Fig. 3C and D).

### Measuring iEEG amplitude modulations as a function of the distance from the calcarine sulcus

A given iEEG measure at each electrode site was interpolated to analysis meshes within 10 mm from the electrode center.^72^ We plotted alpha and high-gamma amplitude modulations as a function of the time bin and 10-mm distance from the calcarine sulcus on the FreeSurfer-flattened cortical surface (Fig. 4B). For example, “Ventral_0-10mm_” denotes the cortical areas within 10 mm from the calcarine sulcus and below the lateral sulcus, whereas “Dorsal_30-40mm_” denotes the cortical areas within 30-40 mm from the calcarine sulcus and above the lateral sulcus. We treated consecutive ≥ eight time-bins showing the 99.99% CI of given iEEG amplitudes beyond the baseline as significant.

**Figure 4.**
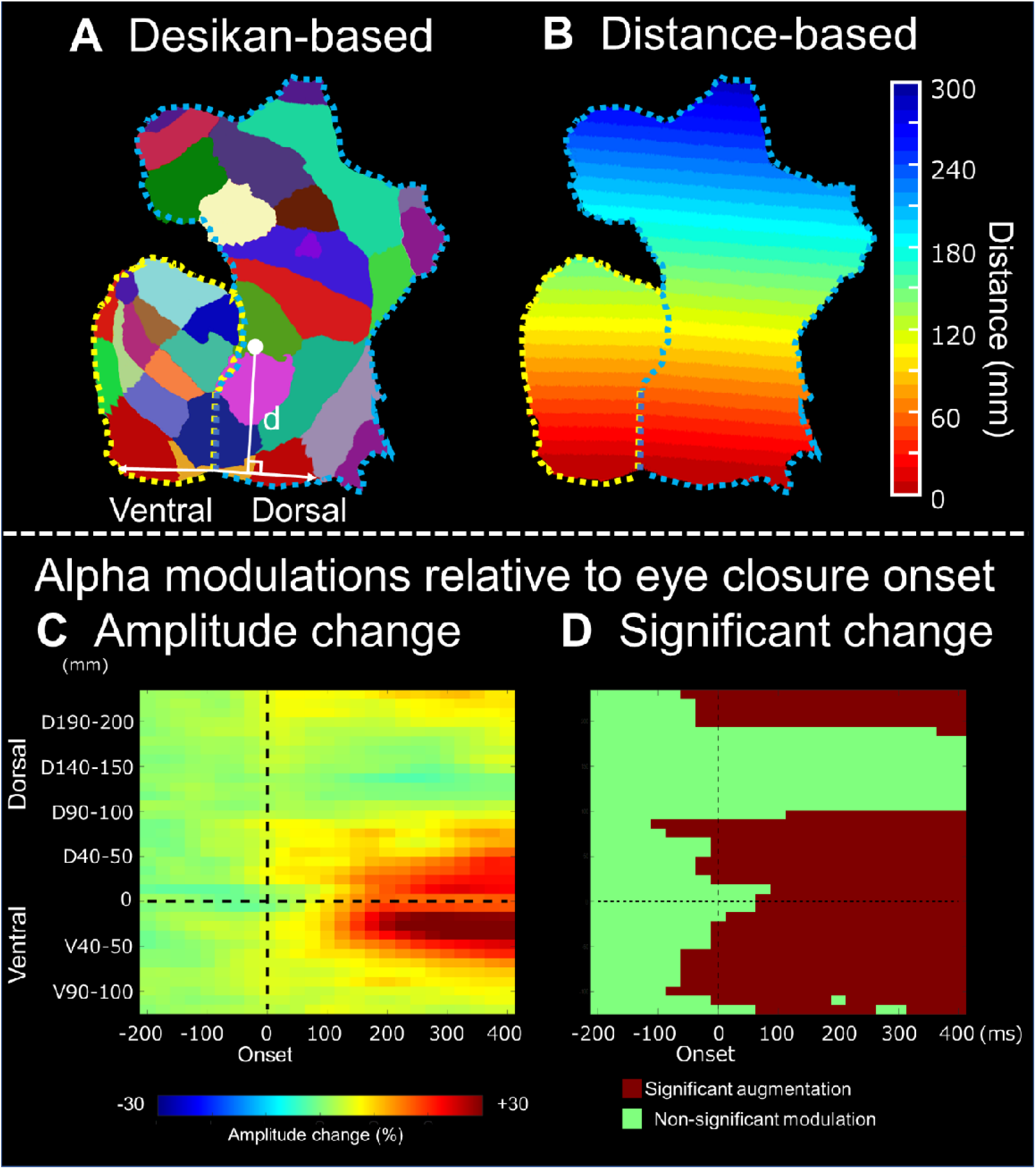
Eye closure-related alpha modulations as a function of the distance from the calcarine sulcus. (A) Each Desikan anatomical region of interest (ROI) is coded with a different color in the flattened cortical map (https://freesurfer.net/fswiki/FreeSurferOccipitalFlattenedPatch). Those inferior to the lateral sulcus were treated as ventral structures (areas surrounded by a yellow dotted line), whereas those superior were treated as dorsal structures (areas surrounded by a blue dotted line) in the distance-based ROI analysis. Refer to Fig. 1B for the meaning of each color code. (B) Each distance-based ROI is color-coded according to the distance from the calcarine sulcus. (C) The percent changes of alpha amplitude modulations are presented as a function of time and distance from the calcarine sulcus (0 ms = eye closure onset). (D) The timing and locations of significant alpha augmentations are denoted as dark red. Supplementary Video 2 presents the alpha amplitude plots at all distance-based ROIs analyzed in the present study.

[**Aim 1**] This distance-based ROI analysis effectively determined whether alpha amplitude augmentation following eye closure would initially occur at the occipital lobe sites more proximal to the calcarine sulcus and subsequently involve more distant cortices (Fig. 4C and D).

### Animation of iEEG amplitude modulations

We sequentially presented alpha and high-gamma amplitude modulations at given electrode sites on the average FreeSurfer pial surface image as a function of time-bin (Fig. 5).^64^ Averaging all patients’ data finally yielded the whole-brain level atlases animating the dynamics of alpha and high-gamma modulations related to eye closure (Supplementary Video 3) and eye opening (Supplementary Video 4).

**Figure 5.**
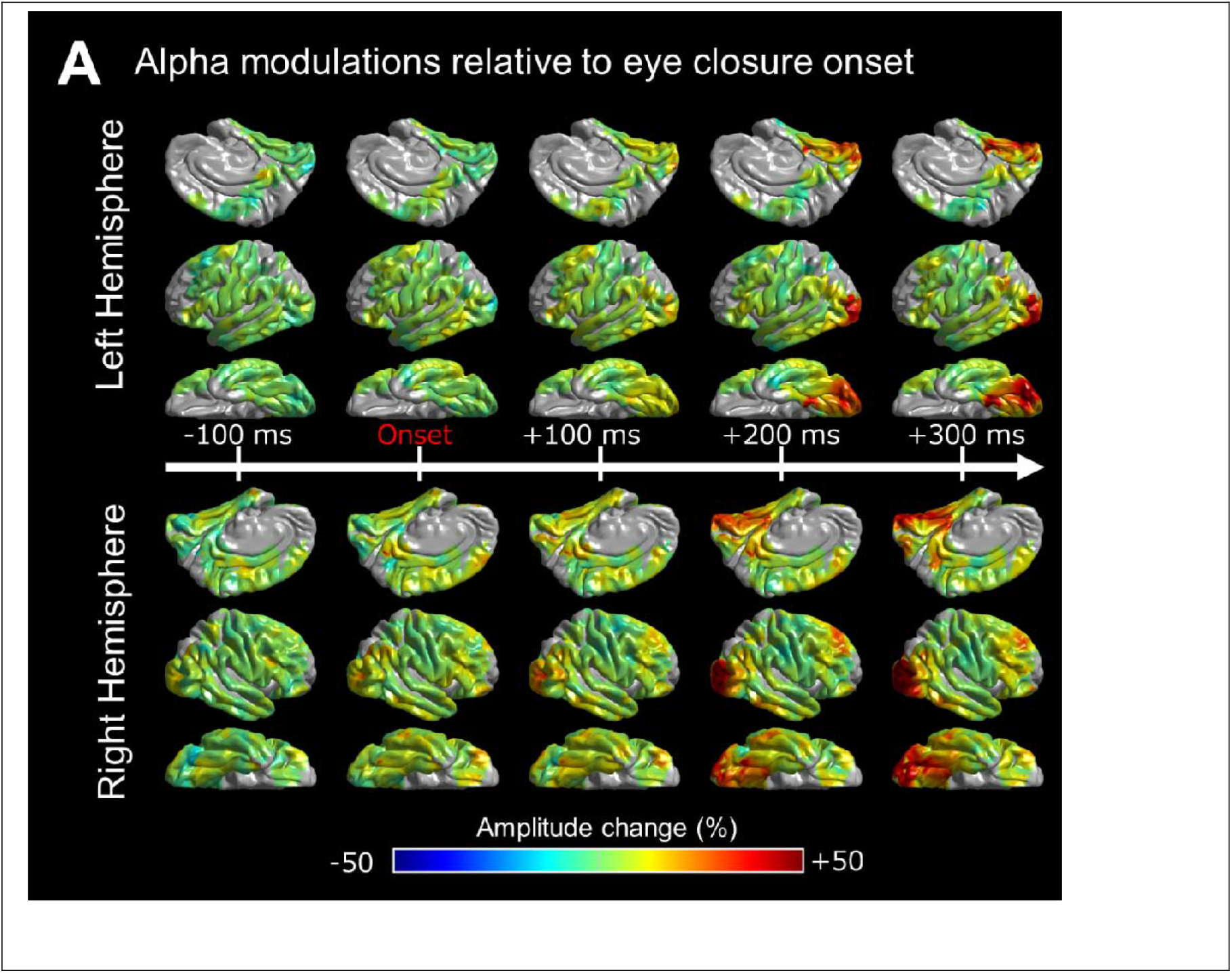
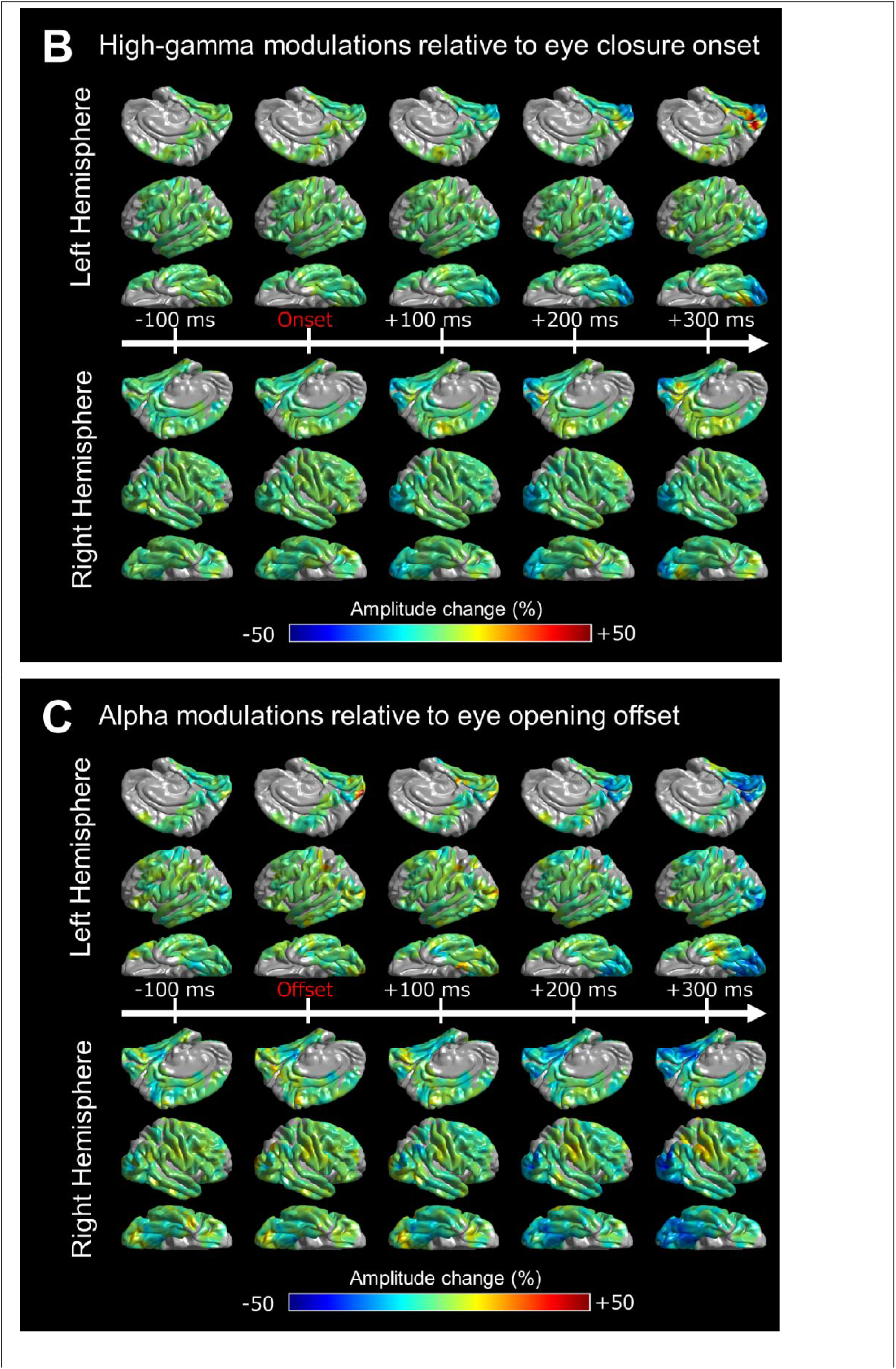
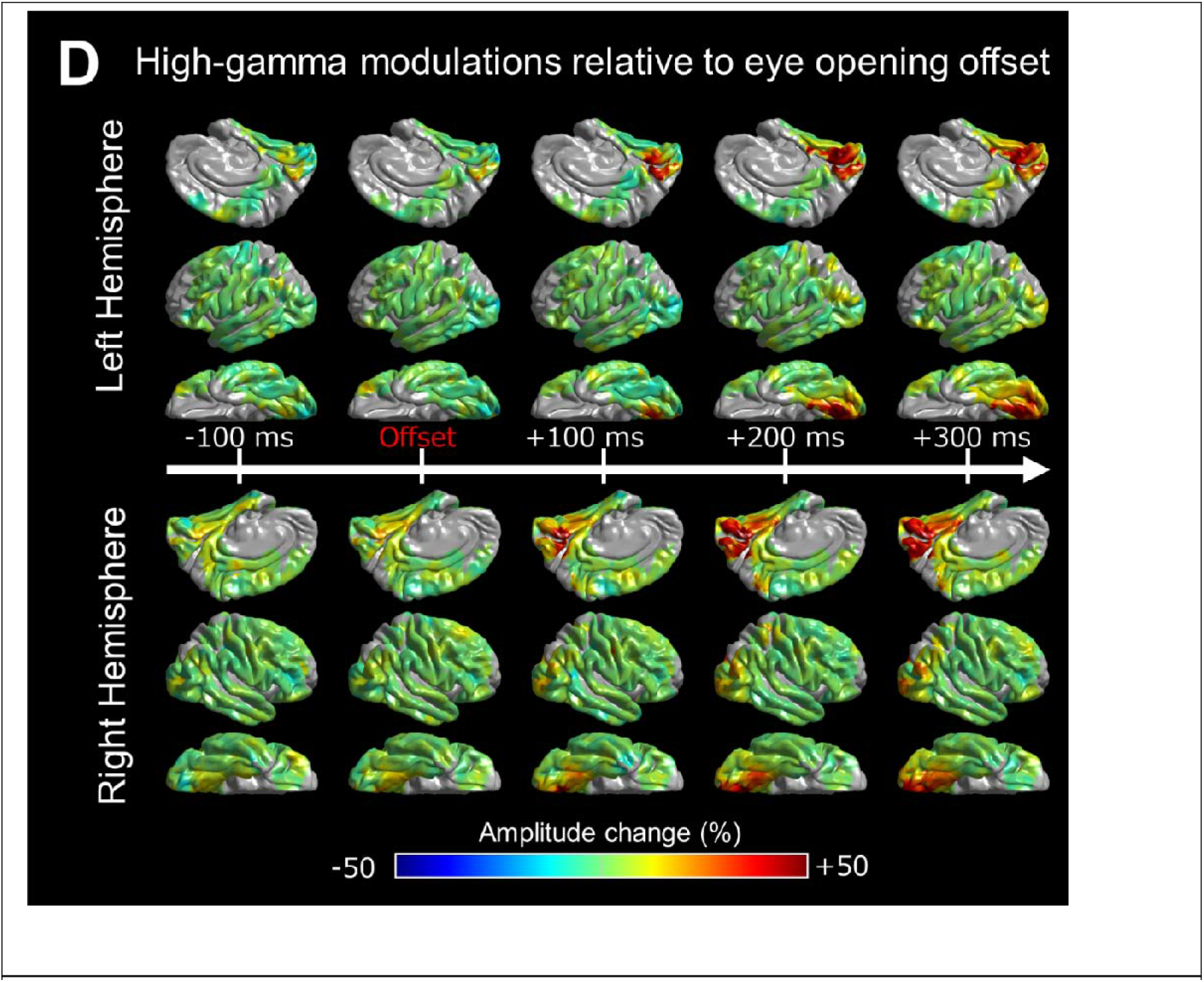
Eye movement-related alpha and high-gamma modulations. The percent change of intracranial EEG amplitude modulations is presented on the FreeSurfer standard brain surface image as a function of time. Eye closure-related (A) alpha and (B) high-gamma modulations (0 ms = eye closure onset; Supplementary Video 3). Eye opening-related (C) alpha and (D) high-gamma modulations (0 ms = eye opening offset; Supplementary Video 4).

### Assessment of functional connectivity dynamics based on alpha co-modulation

With DWI tractography data, we visualized the dynamics of direct white matter functional connectivity between Desikan anatomical ROIs *<u>simultaneously</u>* showing eye closure-related alpha augmentation (or attenuation) lasting at least 200 ms (i.e., ≥ eight 25-ms time-bins; Supplementary Video 5 and Fig. 6). If significant alpha augmentation was noted in χ % of the 2,200-ms analysis time window (i.e., 89 analysis time-bins) on average across 52 ROIs, the chance probability of alpha co-augmentation ≥ 200 ms was ≈ (52/2) × (52-1) × (89-8+1) × {(χ / 100) × (χ / 100)}^8^. With χ of 25%, 30%, and 35%, the estimated chance probability of observing such significant alpha co-augmentation (i.e., Type I error) would be ≈ 0.00003, 0.0005, and 0.006, respectively.

**Figure 6.**
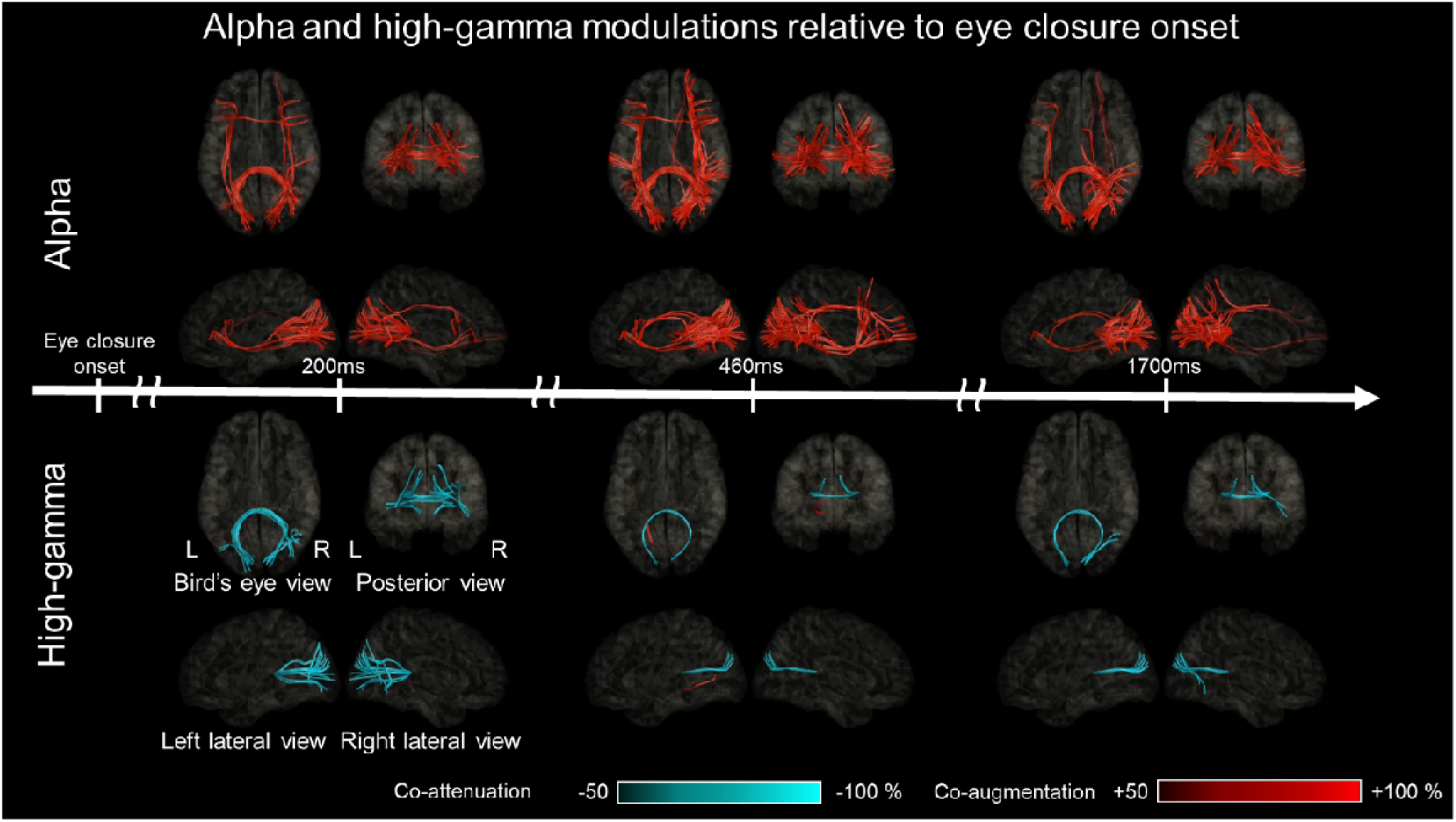
White matter functional connectivity dynamics after eye closure onset. The video snapshots demonstrate the dynamics of functional connectivity alteration between given region pairs occurring after eye closure onset (Supplementary Video 5). Fully-opaque streamlines reflect significant connectivity alterations based on significant alpha co-modulations lasting ≥200-ms (upper images) or high-gamma co-modulations lasting ≥40-ms (lower images). Red streamlines: strengthened functional connectivity. Blue streamlines: weakened functional connectivity.

As previously performed,^42,73^ we used the open-source DWI tractography data from 1,065 individuals participating in the Human Connectome Project (http://brain.labsolver.org/diffusion-mri-templates/hcp-842-hcp-1021).^74^ Our previous study validated the use of open-source DWI data by demonstrating that the dynamics of neural propagations via the white matter pathways based on the open-source data were similar to those found in the individual DWI data.^42^ We placed seeds at ROIs revealing significant alpha co-augmentation lasting ≥200 ms. DSI Studio (http://dsi-studio.labsolver.org/) visualized white matter streamlines directly connecting these ROIs within the Montreal Neurological Institute (MNI) standard space. We considered anatomical DWI streamlines satisfying the following criteria to be legitimate and thus existing: a quantitative anisotropy threshold of 0.05, a maximum turning angle of 70°, a step size of 0 mm, and a streamline length of 10 to 250 mm within the brain parenchyma but outside the brainstem, basal ganglia, and thalamus. Alpha band functional connectivity between a given pair of ROIs at a given 200-ms period was declared to be significantly strengthened (or weakened) if (i) both ROIs showed significant alpha co-augmentation (or attenuation) continuously during the 200-ms period and (ii) legitimate underlying white matter streamlines existed between these ROIs. We assessed alpha-band functional connectivity alteration at each 200-ms epoch in 25-ms sliding windows. **[Aim 3]** Using the dynamic tractography video atlas, we determined whether alpha-based functional connectivity alteration would initially involve the lower-order visual areas and subsequently higher-order areas following eye closure.

We likewise built a dynamic atlas animating the spatiotemporal dynamics of eye opening-related functional connectivity alteration using alpha co-modulation at given pairs of ROIs (Supplementary Video 6 and Fig. 7).

**Figure 7.**
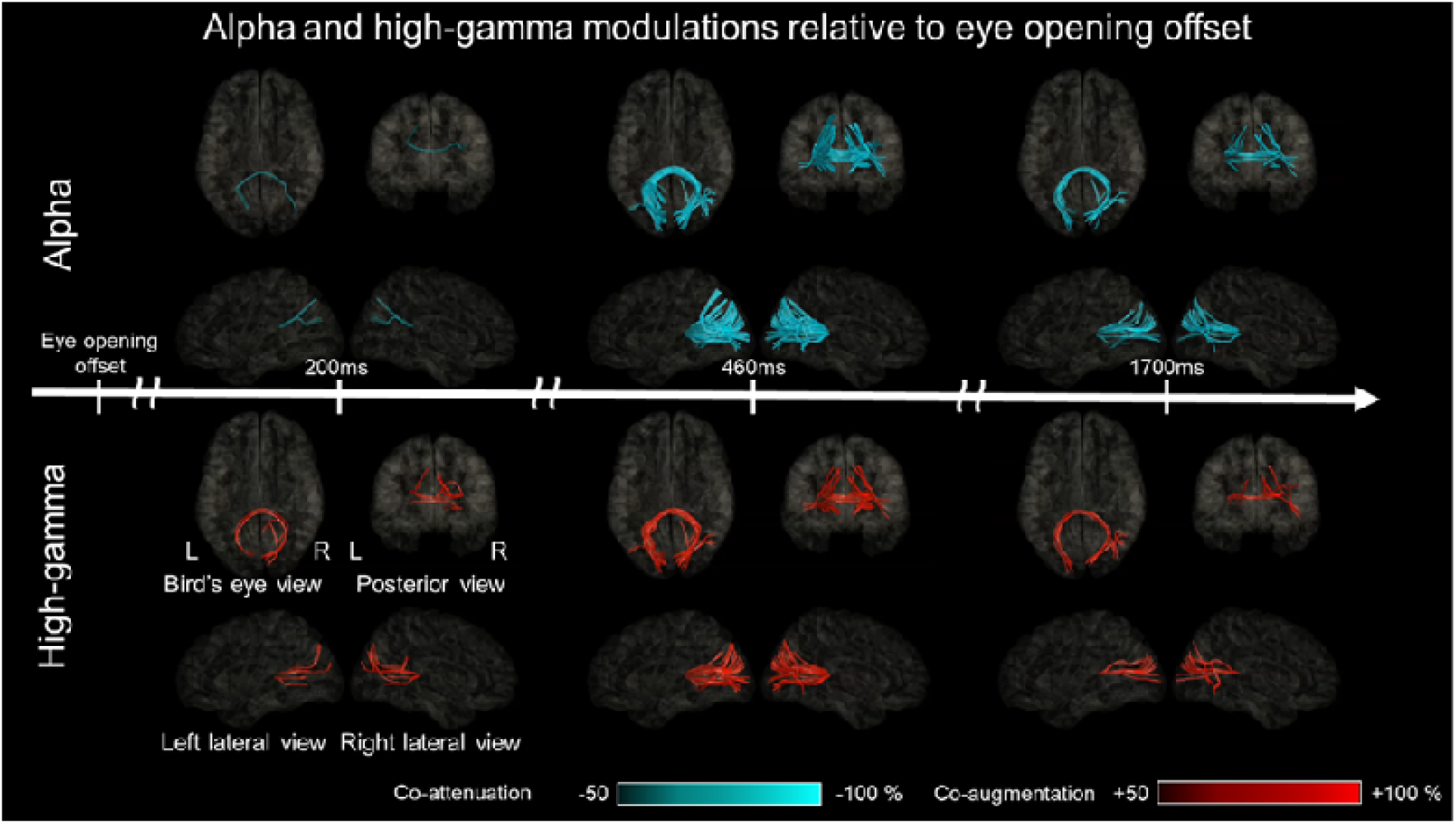
White matter functional connectivity dynamics after eye opening offset. The video snapshots demonstrate the dynamics of functional connectivity alteration between given region pairs occurring after eye opening offset (Supplementary Video 6). Upper: alpha-based connectivity. Lower: high gamma-based connectivity.

### Assessment of functional connectivity dynamics based on high gamma co-modulation

We generated a video atlas presenting the dynamics of eye closure-related alterations of functional connectivity using the streamlines connecting a given pair of ROIs with significant high-gamma co-modulation (i.e., augmentation or attenuation) lasting ≥40 ms (≥ eight 5-ms time-bins; Supplementary Video 5 and Fig. 6). If 10%, 15%, and 20% of the 2,200-ms analysis time window (i.e., 441 analysis time-bins) showed significant high-gamma augmentation on average across 52 ROIs, the estimated chance probability (Type I error) of high-gamma augmentation simultaneously occurring at ≥two ROIs for ≥40 ms was ≈ 6 × 10^−11^, 4 × 10^−8^, and 4 × 10^−8^, respectively.

We likewise built a video atlas animating the dynamics of eye opening-related functional connectivity alteration using high gamma co-modulation at a given pair of ROIs (Supplementary Video 6 and Fig. 7).

### Data/code availability

All data and codes are available upon request to the corresponding author.

## Results

### Behavioral results

The median duration of upward eye movement related to eye closure events was 176 ms across patients (interquartile range [IQR]: 143-217 ms). The duration of downward eye movement related to eye opening events was 175 ms on average across patients (IQR: 128-215 ms).

### [Aim 1] Eye closure-related cortical alpha modulations

The Desikan anatomical ROI-based analysis demonstrated that, at least 50 ms *<u>before</u>* eye closure onset, significant alpha augmentation occurred in multiple ROIs within the occipital and frontal lobes (Supplementary Video 1 and Fig. 3A). We refer to such alpha augmentation as *<u>proactive</u>* because the onset preceded the eye closure onset by more than 39.4 ms (i.e., the temporal resolution of our time-frequency analysis). Expressly, proactive alpha augmentation involved the right lateral occipital (onset latency: -75 ms relative to eye closure onset), right anterior middle frontal gyrus (aMFG) (onset latency: -50 ms), and left posterior inferior frontal gyrus (pIFG) (onset latency: -50 ms) regions. No significant alpha modulation was noted in the frontal eye fields (FEFs) before or after eye closure onset.

Within 100 ms *<u>after</u>* eye closure onset, *<u>reactive</u>* alpha waves (i.e., alpha augmentation after eye closure onset) simultaneously involved widespread brain regions, including lower-and higher-order visual areas as well as those outside the visual areas. Among the lower-order visual areas, lateral occipital regions showed the earliest significant alpha augmentation. The onset latency of significant alpha augmentation (relative to eye closure onset) was +50 ms and -75 ms in the left and right lateral occipital regions, but it was +100 ms and +150 ms in the posterior striatal and +100 ms and +100 ms in the anterior striatal regions, respectively. The maximum degree of alpha amplitude augmentation reached +53.8% and 50.4% in the left and right lateral occipital, +34.0% and +53.5% in the posterior striatal, and +28.1% and +30.2% in the anterior striatal regions.

Eye closure-related alpha augmentation in the lateral occipital region of a given hemisphere preceded that in the higher-order visual areas (Fig. 3A). The onset latency relative to eye closure onset was +75 ms and +175 ms in the left and right posterior fusiform, +350 ms and +75 ms in the pITG, and +150 ms and -25 ms in the inferior parietal regions, respectively. The degree of alpha amplitude augmentation in the higher-order visual areas was smaller than that in the lateral occipital region (Fig. 3A). The maximum degree of alpha augmentation reached +38.1% and 44.1% in the posterior fusiform, +21.0% and +25.0% in the pITG, and +21.9% and +27.1% in the inferior parietal regions, respectively.

Subsets of non-visual areas showed significant eye closure-related alpha augmentation. The onset latency relative to eye closure onset was -50 ms and -25 ms in the bilateral pIFG, -50 ms in the right aMFG, +125 ms and +300 ms in the bilateral posterior middle temporal gyrus (pMTG), and +225 ms and +75 ms in the bilateral posterior superior temporal gyrus (pSTG). The degree of alpha amplitude augmentation in these non-visual areas was smaller than those in the lower- or higher-order visual areas. The maximum degree of alpha amplitude augmentation reached +8.7% and 13.1% in the pIFG, +12.3% in the right aMFG, +13.2% and +18.7% in the pMTG, and +10.2% and +9.3% in the pSTG.

The distance-based ROI analysis also indicated that eye closure-related alpha augmentation rapidly and simultaneously involved widespread brain regions, including lower-order and higher-order visual areas, as well as those outside the visual regions (Supplementary Video 2 and Fig. 4). The earliest significant alpha augmentation occurred in the Dorsal_80-90 ms_ at -100 ms relative to eye closure onset (Fig. 4D), though the degree of amplitude augmentation was modest (Fig. 4C). The degree of alpha amplitude augmentation was high within 50 mm from the calcarine sulcus; the maximum alpha augmentation (+53.1%) was noted for Ventral_20-30 ms_ at +1,425 ms. Dorsal_100-180 mm_, mostly consisting of the pre- and post-central gyri, did not show significant eye closure-related alpha augmentation (Fig. 4C and D).

### [Aim 2] Eye closure-related cortical high-gamma modulations

The Desikan anatomical ROI-based analysis demonstrated that significant high-gamma attenuation simultaneously involved bilateral lower- and higher-order ventral visual areas after eye closure onset (Fig. 3B). The spatial extent and duration of significant high-gamma attenuation were smaller than alpha augmentation (Supplementary Videos 1 and 3). The onset latency of significant high-gamma attenuation was +40 ms and +15 ms in the left and right lateral occipital, +5 ms and +65 ms in the posterior striatal, and +110 ms and +175 ms in the posterior fusiform regions. The maximum degree of high-gamma amplitude attenuation reached -16.1% and -18.0% in the lateral occipital, -19.8% and -20.4% in the posterior striatal, and -8.9% and -9.6% in the posterior fusiform regions. High-gamma attenuation in the lower- and higher-order ventral visual areas became significant after eye closure onset but before eye closure offset (Supplementary Videos 1 and 3).

Significant high-gamma augmentation simultaneously involved the left anterior striatal and parahippocampal regions for 40 ms between +440 ms and +475 ms after eye closure onset; the maximum degree of high-gamma augmentation reached up to +31.0% in the left anterior striatal and +30.0% in the left parahippocampal region. iEEG analysis time-locked to eye closure offset revealed that high-gamma augmentation in the left anterior striatal and parahippocampal regions became significant after eye closure offset (Supplementary Video 1).

Eye closure-related alpha augmentation generally co-occurred with high-gamma attenuation, except in the left anterior striatal region (Fig. 3A and B and Supplementary Video 3). The one-sample t-test revealed that the alpha-by-high-gamma multiplied value was below zero at 24 out of the 28 ROIs showing significant alpha augmentation but above zero at the left anterior striatal region (Bonferroni corrected p<0.05 in each of these 25 ROIs).

### [Aim 3] Functional connectivity dynamics based on eye closure-related alpha co-modulations

The aforementioned time-frequency analysis revealed significant alpha augmentation in 31.6% of the 2,200-ms analysis time window, on average, across 52 ROIs. The estimated Type I error showing a strengthened alpha-based functional connectivity at a pair of two ROIs or more was ≈ 0.0011. The dynamic tractography atlas visualized a strengthening of alpha-based functional connectivity within 125 ms after eye closure onset in the right lateral occipital-pIFG via the inferior fronto-occipital fasciculus (Supplementary Video 5 and Fig. 6). Within 500 ms after eye closure onset, alpha-based functional connectivity was significantly strengthened between homotopic lower- and higher-order visual areas via the posterior corpus callosum, as well as intrahemispheric networks from lower-order visual areas to other lobes (Fig. 6). At 500-1,900 ms after eye closure onset, alpha-based functional connectivity was strengthened in the right medial parietal-frontal networks involving the precuneus, posterior cingulate cortex, and superior frontal gyrus (SFG) via the superior longitudinal fasciculus (Fig. 6).

### [Aim 3] Functional connectivity dynamics based on eye closure-related high-gamma co-modulations

Time-frequency analysis revealed significant high-gamma attenuation and augmentation in 6.8% and 0.1% of the 2,200-ms analysis time window, respectively. The estimated Type I error showing weakened and strengthened high gamma functional connectivity was ≈ 1.3 × 10^−13^ and 1.9 × 10^−40^, respectively. The dynamic tractography atlas visualized a sustained weakening of intra- and inter-hemispheric high gamma-based functional connectivity across the lateral occipital and posterior striatal regions; the weakening of functional connectivity became significant at 90 ms after eye closure onset. Short fibers supported the intra-hemispheric connectivity, whereas the posterior corpus callosum supported the inter-hemispheric connectivity (Supplementary Video 5 and Fig. 6). Conversely, the atlas showed a transient strengthening of the high gamma-based functional connectivity via short fibers between the left anterior striatal and parahippocampal regions at +465 ms.

### [Aim 1] Eye opening-related cortical alpha modulations

The Desikan anatomical ROI-based analysis demonstrated that eye opening-related alpha modulations occurred after eye opening *<u>offset</u>* (Supplementary Video 1 and Fig. 3C). Significant alpha modulation was not noted in the FEFs before or after the eye opening onset. The spatial extent of eye opening-related alpha attenuation was smaller than eye closure-related alpha augmentation. Significant reactive alpha attenuation involved the bilateral lower- and higher-order visual areas, including the lateral occipital (onset latency relative to eye opening *<u>offset</u>*: +200 ms and +175 ms), posterior striatal (+275 ms and +175 ms), anterior striatal (+200 ms and +175 ms), and posterior fusiform regions (+300 ms and +225 ms).

### [Aim 2] Eye opening-related cortical high-gamma modulations

The Desikan ROI-based analysis demonstrated that eye opening-related high-gamma modulations occurred after eye opening *<u>offset</u>* (Supplementary Video 1 and Fig. 3D). Significant high-gamma augmentation initially involved the bilateral lower- and subsequently higher-order visual areas. The onset latency relative to eye opening offset was +75 ms and +10 ms in the anterior striatal, +200 ms and +80 ms in the posterior striatal, +235 ms and +225 ms in the lateral occipital, and +275 ms and +200 ms in the posterior fusiform regions.

Eye opening-related alpha attenuation generally co-occurred with high-gamma augmentation (Supplementary Video 4). The one-sample t-test revealed that the alpha-by-high-gamma multiplied value was below zero at 14 of 16 ROIs that showed significant eye opening-related alpha attenuation (Bonferroni corrected p<0.05 in each of these 14 ROIs).

### [Aim 3] Functional connectivity dynamics based on eye opening-related alpha co-modulations

Significant eye opening-related alpha attenuation occurred in 16.4% of the 2,200-ms analysis time window, on average across 52 ROIs; the estimated Type 1 error showing weakened alpha-based functional connectivity was ≈ 3.1 × 10^−8^. The dynamic atlas visualized a sustained weakening of the intra- and inter-hemispheric alpha-based functional connectivity across lower- and higher-order visual areas, including the lateral occipital, anterior striatal, posterior striatal, and posterior fusiform areas (Supplementary Video 6 and Fig. 7). Short fibers supported most of the intra-hemispheric connectivity pathways, whereas the posterior corpus callosum supported inter-hemispheric connectivity.

### [Aim 3] Functional connectivity dynamics based on eye opening-related high-gamma co-modulations

Significant eye opening-related high-gamma augmentation occurred in 9.2% of the 2,200-ms analysis time window on average across 52 ROIs; the estimated Type I error showing a strengthening of high gamma-based functional connectivity was ≈ 1.6 × 10^−11^. The dynamic atlas visualized a sustained strengthening of the intra- and inter-hemispheric high gamma-based functional connectivity across lower- and higher-order visual areas, including the lateral occipital, anterior striatal, posterior striatal, and posterior fusiform areas (Supplementary Video 6 and Fig. 7). Short fibers supported most of the intra-hemispheric connectivity pathways, whereas the posterior corpus callosum supported the inter-hemispheric connectivity.

## Discussion

### Innovation

The technological innovation of the present study is characterized by animating the rapid alterations of iEEG-based functional connectivity via direct white matter pathways (Fig. 6 and 7 and Supplementary Videos 5 and 6). The high gamma-based atlas animated the connectivity pathways at given 40-ms epochs using a 5-ms sliding window, while the alpha-based one used 200-ms epochs with a 25-ms sliding window. Direct iEEG signal sampling from deeply-seated areas allowed us to segregate between the contrasting high-gamma dynamics involving the posterior and anterior striatal regions, which is not feasible with noninvasive recording (Supplementary Videos 3 and 4). We declared that direct functional connectivity was significantly altered only when [i] iEEG amplitude modulations at a pair of regions were temporally coupled and [ii] biologically-plausible, direct white matter pathways were present on DWI tractography.

The dynamic tractography atlas, based on our novel iEEG-DWI multimodality analysis, revealed eye closure-related alterations of intra- and inter-hemispheric functional connectivity (Supplementary Video 5). For example, significant and sustained alpha augmentation simultaneously involved the lateral occipital and pIFG regions in each hemisphere, within 500 ms after eye closure onset (Fig. 3A). The results also suggest that the inferior fronto-occipital fasciculus supports intra-hemispheric functional connectivity alteration, whereas the posterior corpus callosum supports the inter-hemispheric connectivity. Although the lateral occipital and pIFG simultaneously showed significant eye closure-related alpha augmentation, our dynamic tractography indicated the absence of inter-hemispheric connectivity between these regions due to a lack of legitimate direct streamlines. We expect that the video materials will be beneficial to better understand the network dynamics of electrocorticographic reactivity to eye movements. The normative dynamic atlas can be an asset for clinicians to understand the significance of scalp EEG alpha waves while assessing the functional integrity of the cerebrum. Our atlas is also expected to help understand the significance of task-related alpha modulations in cognitive neuroscience investigations. Our results warrant further study to consider if given alpha modulations (e.g., at pIFG) can be attributed to different eye movement patterns between task and baseline periods.

### Significance of functional connectivity modulations

Increased functional connectivity strength based on *<u>high-gamma</u>* co-augmentation likely reflects network activation via a given white matter pathway (Fig. 7), whereas high-gamma co-attenuation reflects network deactivation (Fig. 6). Our previous iEEG study demonstrated that direct effective connectivity was strengthened across cortical areas simultaneously showing task-related high-gamma augmentation; thereby, the connectivity strength was rated by early cortico-cortical spectral responses induced by local single-pulse electrical stimulations.^42^ Investigators have suggested that neuronal circuits simultaneously engaging in high-frequency activities can develop use-dependent, direct functional connectivity.^40,41^ Resection of cortical areas showing task-related high-gamma activity was found to increase the risk of postoperative cognitive decline.^36^

In contrast, functional connectivity alterations based on alpha co-augmentation need to be interpreted with caution. Investigators suggest that eye closure-related alpha augmentation is correlated to functional idling of cortical areas.^2,3,37^ Task-related alpha attenuation has been reported to be temporarily coupled with high-gamma augmentation in many iEEG studies of motor,^15,75,76^ somatosensory,^77,78^ visual,^16,17,79,80^ auditory,^81,82^ memory,^83,84^ and language function.^85-88^ Investigators suggest that task-related high-gamma augmentation is more accurately time-locked to a given behavior and spatially more confined to the eloquent cortex.^15,75-77^ Our previous iEEG study demonstrated that naming-related high-gamma augmentation predicted postoperative cognitive declines better than alpha attenuation.^36^ Thus, investigators may hypothesize that functional connectivity alteration based on alpha co-augmentation reflects increased idling of a given white matter network and a reduced chance of neuronal activation.^89,90^ In turn, alpha co-attenuation might reflect reduced idling and an increased chance of neuronal activation.

### Physiological significance of *<u>proactive</u>* alpha waves

Our novel observations include proactive alpha modulations, which occurred in multiple regions before eye closure onset. Significant alpha augmentation occurred in the right lateral occipital region 75 ms before, right aMFG 50 ms before, and left pIFG 50 ms before eye closure onset. Such proactive alpha co-augmentation can be attributed to patient behavioral status, such as reduced visuospatial attention and internal verbal thought. Previous iEEG studies of humans and non-human primates reported that reduced visuospatial attention was reflected by increased amplitude of alpha activity in the large-scale networks involving visual areas, whereas increased attention was coupled with alpha attenuation.^91-95^ Other iEEG studies reported that alpha amplitude in the left prefrontal regions was higher during the resting period than in the active period within a required linguistic process.^27,36,85,96^

We do not have definitive evidence that proactive alpha augmentation in the aforementioned cortical areas is causally and directly associated with the initiation of eye closure. No significant alpha augmentation was noted in the FEFs before or after eye closure onset. No high-gamma augmentation or attenuation was observed before eye closure onset in any ROI.

### Physiological significance of <u>reactive</u> eye closure-related alpha and high-gamma *activities*

The observed temporal dynamics of alpha amplitude augmentation do not support the notion that eye closure-related alpha augmentation uniformly reflects feedforward or feedback rhythms propagating from lower to higher-order visual cortex, or *vice versa* (Supplementary Videos 3 and 5). Rather, significant alpha augmentation co-occurred in extensive networks within and outside the visual areas within 500 ms after eye closure onset. The bilateral, lateral occipital regions showed the earliest onset latency and the highest degree of alpha augmentation among lower-order visual areas. Reactive alpha augmentation was also noted in the higher-order ventral and dorsal visual areas, including the posterior fusiform, pITG, and inferior parietal regions. Dynamic tractography demonstrated that these lower- and higher-order visual areas had strengthened alpha-based functional connectivity with homotopic areas via the posterior corpus callosum. Furthermore, these visual areas had strengthened alpha functional connectivity with the non-visual areas, including the pIFG in each hemisphere, via the inferior fronto-occipital fasciculus (Supplementary Video 5).

Unlike the extensive distribution of such alpha augmentation, eye closure-related high-gamma attenuation was confined to the lateral occipital, posterior striatal, and posterior fusiform regions; this observation is consistent with a snapshot showing the spatial characteristics of alpha and high-gamma modulations 4-6 seconds after eye closure.^17^ The present study demonstrated that these visual areas showed significant high-gamma attenuation within 175 ms after eye closure onset. Such spatio-temporal coupling between alpha augmentation and high-gamma attenuation is consistent with the notion that reactive alpha augmentation in these visual areas reflects an idling rhythm, and that they were indeed deactivated after eye closure onset, as reflected by high-gamma attenuation.

A notable finding was that significant high-gamma augmentation occurred in the left anterior striatal and parahippocampal regions 440-475 ms after eye closure offset, despite the simultaneous alpha augmentation (Supplementary Video 3). The anterior striatal region, located in the anterior-medial occipital area, is suggested to represent the low-order visual cortex for peripheral visual fields, whereas the posterior striatal region represents parafoveal fields.^97-100^ Thus, reactive eye closure-related high-gamma augmentation in the left anterior striatal region may reflect increased vigilance to surroundings or visual imagination. An fMRI study of healthy adults reported that, in complete darkness, the anterior striatal regions showed increased hemodynamic responses during sustained eye closure than opening.^101^ An alternative explanation for high-gamma augmentation in the left anterior striatal region is neural activation passively elicited by a massive change in the visual input at the retinal level during eye closure offset. The anterior striatal cortex preferentially receives visual inputs from the retinal rod cells, which play a crucial role in detecting light and dark contrast.^102,103^ Interpretation of co-occurring alpha and high-gamma augmentation is not straightforward. A possible explanation for this finding is that the anterior striatal region was capable of generating high-gamma augmentation despite the enhanced idling rhythm, as reflected by alpha augmentation. Investigators have suggested that local neural activation is more closely reflected by high-gamma augmentation than modulations of other frequency bands.^36,104^

### Significance of <u>reactive</u> eye opening-related high-gamma and alpha modulations

Between eye opening onset and offset, visual inputs at the retinal level should have been rapidly altered, but no significant high-gamma augmentation was noted in the visual areas during this time window. The lack of high-gamma augmentation during active eye movement can be attributed to the visual cortex’s active preparation for massive image motion expected during vertical eye movements; this cortical mechanism is referred to as ‘efferent copy’ and has been reported in previous studies of saccade-related iEEG modulations.^62,105,106^

After the *<u>offset</u>* of eye opening, significant high-gamma augmentation initially occurred in the anterior striatal, subsequently in the posterior striatal, and lastly in the lateral occipital and posterior fusiform regions. This observation supports the notion that high-gamma activity is driven by the visual stimuli available at the eye opening offset and that such reactive high-gamma augmentation propagates from lower- to higher-order visual areas. The propagation of high-gamma augmentation mentioned above is, in part, similar to those induced by full-field flash stimuli during eye closure; though, such simple flash stimuli did not elicit high-gamma augmentation in the posterior fusiform regions (see the video file in Nakai et al.^107^). In contrast, the bilateral posterior fusiform regions showed high-gamma augmentation reactive to the complex natural scene presented at the offset of eye opening (Supplementary Video 4 of the present study).

The dynamic tractography atlas visualized a sustained strengthening of intra- and inter-hemispheric high gamma-based functional connectivity across the lower- and higher-order visual areas. The dynamics of intra-hemispheric functional connectivity likely reflect the hierarchical object and place recognition processes initiated at the offset of eye opening.^68-71^ The strengthening of inter-hemispheric functional connectivity via the posterior corpus callosum may contribute to the transformation and integration of visual inputs processed in each hemisphere.^108,109^ This notion is supported by the observation that some patients with drug-resistant epilepsy developed alexia - neglect of the visual field ipsilateral to the language dominant hemisphere - following total corpus callosotomy.^108,110,111^ Our previous iEEG study of three patients showed that single-pulse electrical stimulation of a cortical region elicited a cortico-cortical spectral response in the homotopic area of the contralateral hemisphere in an average of 22 ms.^112^

After the *<u>offset</u>* of eye opening, significant alpha attenuation initially occurred in the anterior striatal regions, subsequently involved the posterior striatal and lateral occipital regions, and finally the posterior fusiform regions. Supplementary Video 4 best demonstrates that eye opening-related alpha attenuation was coupled with high-gamma augmentation in both central and peripheral visual areas. The onset latency of reactive alpha attenuation occurred several hundred milliseconds after high-gamma augmentation. The observed temporal dynamics of high-gamma and alpha modulations were similar to those reported in a previous study of somatosensory-related iEEG amplitude modulations: high-gamma augmentation preceded alpha attenuation by several hundred milliseconds.^77^ These observations are in line with the notion that high-gamma augmentation closely reflects local neural activation.

### Methodological limitations

The present study is not designed to clarify the direction of information transfers. Computational integration of Granger-type iEEG effective connectivity measures,^92,113-118^ iEEG responses to single-pulse electrical stimulation,^42,112^ and streamlines on DWI tractography^42,74^ may be necessary to accurately assess the directionality of direct neural information transfers via given white matter pathways.

In the present study, all patients had iEEG sampling from all four lobes of each hemisphere to determine the boundaries between the epileptogenic zone and eloquent cortices. Yet, limited spatial sampling is an inevitable limitation of any iEEG study. None of our patients had iEEG signal sampling from the thalamic nuclei because there was no clinical need. Thus, our study is not designed to clarify the network dynamics involving the thalamus. Deep brain stimulation of the thalamus has become one of the standard palliative treatments for selected patients with drug-resistant epilepsy, and stereotactic EEG recording from the thalamic nuclei is expected to be a common clinical procedure for localizing the therapy target.^24,119,120^ Future collaborative studies may provide an opportunity to determine the cortico-cortical and thalamocortical network dynamics of iEEG modulations around eye movements.

The number of spontaneous eye movement events detected by video and EOG differed across patients; a patient with 77 eye movement events provided iEEG data with a signal-to-noise ratio twice as good as those with 16 events. Unlike a previous study,^24^ we did not employ an auditory-cued eye closure/opening task. We are also aware that the number of electrode sites differed across ROIs; for example, the right lateral occipital regions had 67 electrode sites, whereas the right FEF had 30; this indicates that the statistical power was 1.5 times better in the right lateral occipital region compared to the right FEF. Thus, failure to find a significant alteration in network dynamics might be attributed to insufficient numbers of trials or electrode sites within given ROIs. However, the variation in the number of trials and iEEG electrode sites is inevitable since we do not expand the duration nor extent of iEEG recording for research purposes. Nonetheless, the signal fidelity of iEEG recording is >100 times better than scalp EEG recording.^14^ Thus, investigators suggest that even a single trial of iEEG signals can provide meaningful information.^29,121,122^

The present study did not include patients younger than five years because the posterior dominant rhythm may range below 8 Hz in healthy children. The inclusion of such very young children is expected to allow us to determine the ontogenic changes in the network dynamics of iEEG modulations before and after eye movements. Such studies may improve the understanding of how the human brain develops network dynamics supporting object recognition, interhemispheric integration of visual inputs, and efferent copy during eye movements.

## Abbreviations

aMFG: anterior middle frontal gyrus
CI: confidence interval
DMNs: default mode networks
DWI: diffusion-weighted imaging
EEG: electroencephalography
EOG: electrooculography
FEFs: Frontal eye fields
FIR: finite impulse response
fMRI: functional magnetic resonance imaging
HFOs: high-frequency oscillations
iEEG: intracranial EEG
IQR: interquartile range
LGN: lateral geniculate nucleus
MEG: magnetoencephalography
MNI: Montreal Neurological Institute
PDR: posterior dominant rhythm
pIFG: posterior inferior frontal gyrus
pITG: posterior inferior temporal gyrus
pMTG: posterior middle temporal gyrus
pSTG: posterior superior temporal gyrus
ROI: region of interest
SOZ: seizure onset zone
3D: three-dimensional.

## Acknowledgments

We are grateful to Karin Halsey, BS, REEGT, and Jamie MacDougall, RN, BSN, CPN at Children’s Hospital of Michigan, for the collaboration and assistance in performing the studies described above.

## CRediT authorship contribution statement

- Hiroya Ono: Conceptualization, Investigation, Data curation, Formal analysis, Visualization, Writing the original draft.
- Masaki Sonoda: Conceptualization, Investigation, Data curation, Formal analysis (including the distance-based region of interest analysis), Methodology, Visualization, Software.
- Kazuki Sakakura: Data curation, Formal analysis (including the dynamic tractography), Methodology, Visualization, Software.
- Yu Kitazawa: Formal analysis, Methodology, Visualization, Software.
- Takumi Mitsuhashi: Methodology, Visualization, Software.
- Ethan Firestone: Revising the manuscript.
- Aimee F. Luat: Data acquisition.
- Neena I. Marupudi: Data acquisition.
- Sandeep Sood: Data acquisition.
- Eishi Asano: Conceptualization, Investigation, Data acquisition, Formal analysis, Methodology, Visualization, Resources, Supervision, Funding acquisition, Writing the original draft, Revising the manuscript.

## Additional Contributions

### Funding

This work was supported by the National Institutes of Health (NS064033 to EA).

