## Supplementary Table 1 for "Dynamic cortical and tractography atlases of proactive and reactive alpha and high-gamma activities"

**in**

This document includes
**Supplementary Table 1**

**Legends for Supplementary Videos 1-6**

**Supplementary Table 1: The number of electrodes at given regions of interest (ROIs).**

| ROIs | Left | Right | Total |
| --- | --- | --- | --- |
| frontal eye field (FEF) | 23 | 30 | 53 |
| anterior fusiform gyrus (aFG) | 10 | 23 | 33 |
| posterior fusiform gyrus (pFG) | 36 | 36 | 72 |
| anterior inferior-frontal gyrus (aIFG) | 22 | 36 | 58 |
| posterior inferior-frontal gyrus (pIFG) | 78 | 84 | 162 |
| inferior parietal lobule (IPL) | 18 | 40 | 58 |
| anterior inferior-temporal gyrus (aITG) | 15 | 22 | 37 |
| posterior inferior-temporal gyrus (pITG) | 23 | 16 | 39 |
| lateral-occipital gyrus (LOG) | 65 | 67 | 132 |
| anterior middle-frontal gyrus (aMFG) | 53 | 96 | 149 |
| posterior middle frontal gyrus (pMFG) | 55 | 81 | 136 |
| anterior middle-temporal gyrus (aMTG) | 22 | 19 | 41 |
| posterior middle-temporal gyrus (pMTG) | 53 | 38 | 91 |
| paracentral gyrus (PCL) | 7 | 11 | 18 |
| postcentral gyrus (PoCG) | 103 | 105 | 208 |
| parahippocampal gyrus (PHG) | 4 | 10 | 14 |
| posterior cingulate gyrus (pCG) | 15 | 22 | 37 |
| precentral gyrus (PreCG) | 72 | 125 | 197 |
| precuneus (PCun) | 6 | 18 | 24 |
| anterior striatal gyrus (aSG) | 34 | 54 | 88 |
| posterior striatal gyrus (pSG) | 20 | 16 | 36 |
| superior frontal gyrus (SFG) | 37 | 71 | 108 |
| superior parietal lobule (SPL) | 16 | 12 | 28 |
| anterior superior-temporal gyrus (aSTG) | 27 | 40 | 67 |
| posterior superior-temporal gyrus (pSTG) | 55 | 59 | 114 |
| supramarginal gyrus (SMG) | 81 | 73 | 154 |
| medial orbitofrontal gyrus (MOrb) | 2 | 1 | 3 |
| temporal pole (TP) | 0 | 2 | 2 |
| entorhinal gyrus (Ent) | 1 | 4 | 5 |
| anterior cingulate gyrus (aCG) | 1 | 5 | 6 |

**Supplementary Table 1. The number of electrodes at regions of interest (ROIs).** The ROI analysis was not performed at the medial orbitofrontal gyrus (MOrb), temporal pole (TP), entorhinal gyrus (Ent), or anterior cingulate gyrus (aCG) because of the limited number of electrode sites eligible for analysis.

**Supplementary Video 1. Eye movement-related alpha and high-gamma modulations.** A given plot presents the percent change in alpha and high-gamma amplitudes at each Desikan-based anatomical region of interest (ROI) compared to the baseline mean at 200 to 600-ms before eye movement onset (Fig. 3). This video initially presents intracranial EEG amplitude modulations relative to eye closure onset and offset, and it subsequently presents those relative to eye opening onset and offset. Solid line: mean across all available electrode sites within a given ROI. Shade: 99.99% confidence interval (Nakai et al., 2019). Magenta: Left hemisphere. Green: Right hemisphere.

[Supplementary reference] Nakai Y, Sugiura A, Brown EC, et al. Four-dimensional functional cortical maps of visual and auditory language: Intracranial recording. Epilepsia 2019;60:255-267.

**Supplementary Video 2. Eye movement-related alpha and high-gamma modulations as a function of the distance from the calcarine sulcus.** A given plot presents the percent change in alpha and high-gamma amplitudes compared to the preceding baseline mean at 200 to 600-ms before eye movement onset at each distance-based region of interest (ROI) (Fig. 4; Sakakura et al., 2022). The video presents intracranial EEG amplitude modulations relative to eye closure onset. Solid line: mean across all available electrode sites within a distance-based ROI. Shade: 99.99% confidence interval. Orange: Dorsal regions. Purple: Ventral regions.

[Supplementary reference] Sakakura K, Sonoda M, Mitsuhashi T, et al. Developmental organization of neural dynamics supporting auditory perception. Neuroimage 2022;258:119342.

**Supplementary Video 3. Eye closure-related cortical alpha and high-gamma modulations.** The dynamic atlas demonstrates the spatiotemporal modulations of alpha and high-gamma modulations relative to the onset and offset of eye closure. Fig. 5A and B show the snapshots of alpha and high-gamma modulations related to eye closure.

**Supplementary Video 4. Eye opening-related cortical alpha and high-gamma modulations.** The dynamic atlas demonstrates the spatiotemporal modulations of alpha and high-gamma modulations relative to the onset and offset of eye opening. Fig. 5C and D show the snapshots of alpha and high-gamma modulations related to eye opening.

**Supplementary Video 5. White matter functional connectivity alterations related to eye closure.** The video demonstrates the dynamics of functional connectivity alteration between given region pairs occurring relative to eye closure onset and offset (Fig. 6). Left: alpha-based dynamic tractography atlas, which visualizes the proportion of alpha co-modulation at each 200-ms epoch in 25-ms sliding windows. Right: high gamma-based dynamic tractography atlas, which visualizes the proportion of high gamma co-modulation at each 40-ms epoch in 5-ms sliding windows. Red streamlines reflect strengthening of functional connectivity, whereas blue ones reflect weakening. Fully-opaque streamlines indicate an occurrence of amplitude co-modulation during all eight time-bins and a significant connectivity alteration during the epoch.

**Supplementary Video 6. White matter functional connectivity dynamics related to eye opening.** The video demonstrates the dynamics of functional connectivity alteration between given region pairs occurring relative to eye opening onset and offset (Fig. 7). Left: alpha-based dynamic tractography atlas, which visualizes the proportion of alpha co-modulation at each 200-ms epoch in 25-ms sliding windows. Right: high gamma-based dynamic tractography atlas, which visualizes the proportion of high gamma co-modulation at each 40-ms epoch in 5-ms sliding windows. Red streamlines reflect strengthening of functional connectivity, whereas blue ones reflect weakening. Fully-opaque streamlines indicate an occurrence of amplitude co-modulation during all eight time-bins and a significant connectivity alteration during the epoch.
